# smallrnaseq: short non coding RNA-Seq analysis with Python

**DOI:** 10.1101/110585

**Authors:** Damien Farrell

## Abstract

The use of next generation sequencing is now a standard approach to elucidate the small non-coding RNA species (sncRNAs) present in tissue and biofluid samples. This has revealed the wide variety of RNAs with regulatory functions the best studied of which are microRNAs. Profiling of sncRNAs by deep sequencing allows measures of absolute abundance and for the discovery of novel species that have eluded previous methods. Specific considerations must be made when quantifying and cataloging sncRNAs and multiple algorithms are now available, mostly focused on miRNA analysis. *smallrnaseq* is a Python package that implements some of the standard approaches for quantification and analysis of sncRNAs. This includes miRNA quantification and novel miRNA prediction. A command line interface makes the software accessible for general users.

## Introduction

Small RNA is a term used rather broadly to describe short regulatory non coding RNAs found in eukaryotes. This may include longer molecules such as transfer RNAs and snRNAs. However the term most often refers to the very short RNAs of about 20–30 nucleotides long that are important in the RNA silencing pathway by their association with Argonaute (Ago)-family proteins. These are classified based on their biogenesis mechanism and the type of Ago protein that they are associated with. microRNAs (miRNAs), endogenous small interfering RNAs (endo-siRNA) and Piwi-interacting RNAs (piRNAs) are the predominant classes that have been identified. Though recent discoveries of non-canonical small RNAs have blurred the boundaries between the classes^1^.

Advances in next generation sequencing technologies in the past decade have led to intensive discovery in this field. Many studies have now been done in cataloging miRNA genes in particular and establishing databases such as miRBase ^2^. Most of the abundant miRNAs have likely been discovered in model organisms like humans, mice and other mammalian species. Identifying less abundant and/or cell type-specific ncRNAs and their biological relevance is still an ongoing task.

Protocols have been developed that enrich for miRNAs and piRNAs in an effort to exclude RNA turnover and hydrolysis products arising from rRNAs and tRNAs^3^. Library preparation involves consecutive steps of adapter oligonucleotide RNA ligation. Adapter-ligated RNA is then size-fractionated on polyacrylamide gels with the band in the appropriate size range excised. Finally, reverse transcription and PCR are performed and the resulting small cDNA library is deep sequenced. The Illumina platform commonly used typically yields tens of millions of reads almost always 50 bp in length with multiplexing allowing 20 or more samples to be run at once. The resulting reads are returned in the form of fastq files which must be analyzed to yield expression profiles of known and novel genes. sncRNA datasets have several commonly seen features. Firstly, the abundance distribution is highly skewed with a few genes constituting a majority of the total reads. Secondly, especially in the case of miRNAs, a read may map to multiple precursors and/or possibly other locations in the genome. This has an effect on both alignment parameters and the method used to calculate final read counts. Finally, there is a distribution of isoforms (called isomiRs for miRNAs) that display sequence variations, typically by changes at the 3’ end. Often the most abundant miRNA differs from the miRBase sequence, probably because the dominant isomiR varies by tissue and sample. This has an important consequence for the use of miRNA based markers since in certain commercial assays the wrong isoform may be used as the sensor^4^. The length of the reads means that a good deal of adapter dimer may be seen in the data returned from a sequencing run, depending on library preparation.

Analyzing small RNA sequencing data remains a challenging process not only because of the technical difficulties mentioned above. There is also a practical problem of appropriate software selection and usage. Though multiple tools^5^ have been created only a subset are user friendly enough for general users. The papers describing these tools are sometimes highly technical and confusing for those not experienced in bioinformatic methods. Most are focused on miRNA profiling^6^–^9^ of which the most well known is probably miRDeep2^10^. Some tools are web-based making them convenient to access but this usually requires uploading data to a server which is not suitable for a large scale study with many large sample files.

This article details a Python package called *smallrnaseq* that aims to simplify and de-mystify this type of analysis whilst still allowing enough extensibility for more advanced users. It provides an independent implementation of some previously described algorithms such as isomiR labelling and novel miRNA prediction. The main aims of this software are:

- provide a relatively simple but flexible command line tool
- allow general analysis of small RNAs and not just miRNAs
- provide a Python API for programmers who want to integrate the modules into their own custom work flow or as part of another tool (see discussion)

## Results

Small RNA seq analysis typically follows the steps shown in Figure 1. All of these steps are described below. Trimming adapters should always be done in advance of processing unless using an aligner that allows soft clipping of reads (not Bowtie). This can be done with a tool like cutadapt.

**Figure 1.**
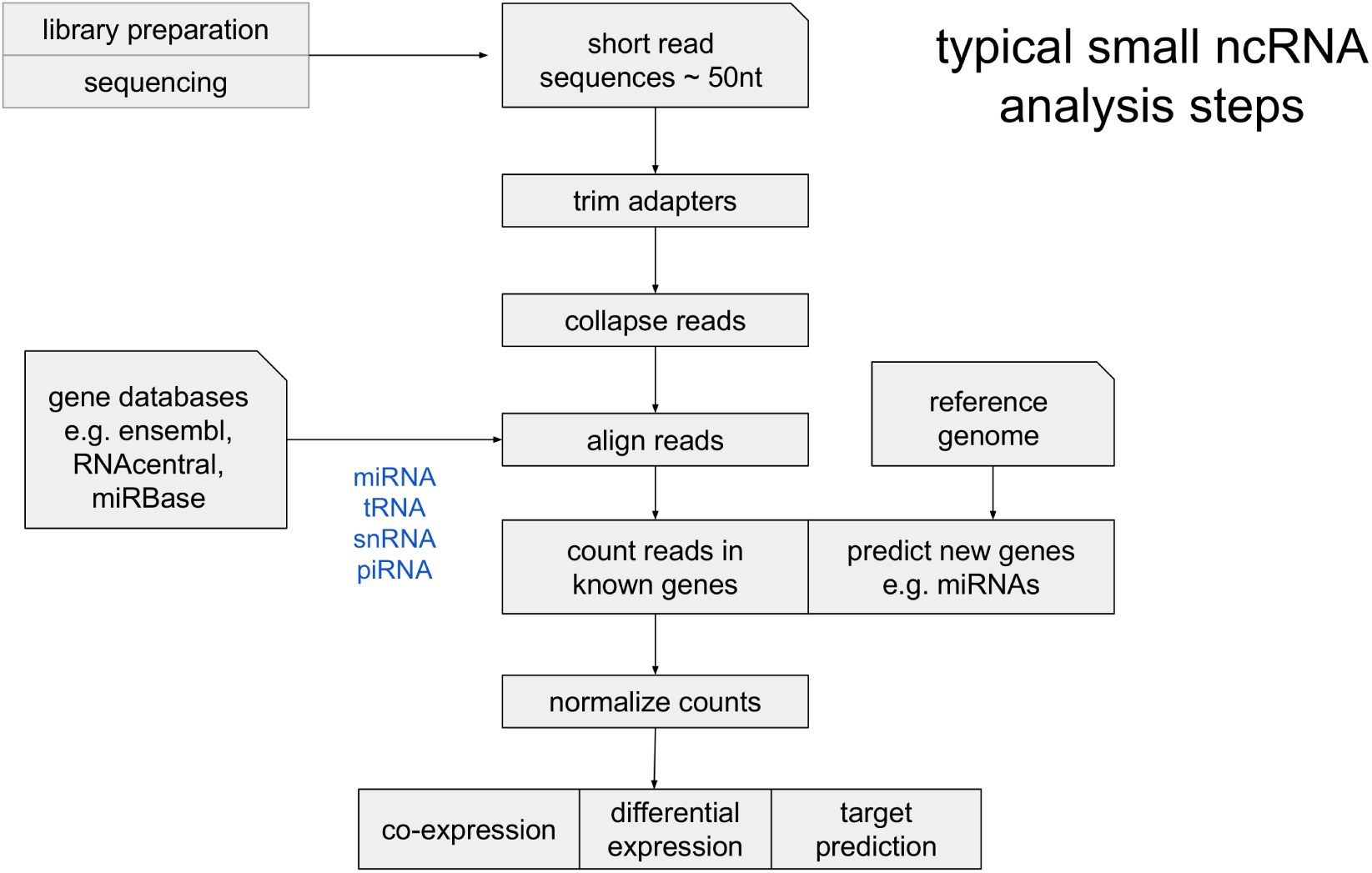
Typical workflow for sncRNA seq analysis. This is almost the same as for standard RNA-seq except that the reads are 50 nt in length which affects alignment considerations.

### Command line interface

Installing the package provides a command ’smallrnaseq’ that is a command line interface to the library without the need for any Python coding. Usage simply requires entering options into a text configuration file and having the proper input files prepared. The advantage of configuration files is that they avoid long commands with many options having to be typed, which is prone to mistakes. Also the files can be stored to recall what setting was used, to copy them for another data set or to share with other users. The meaning of each option is explained in the documentation in detail. Functionality currently available from the command line is explained in the following sections. This includes counting of any user supplied RNA gene annotations, miRNA quantification, novel miRNA prediction and counting of genomic features. The algorithm is designed to handle multiple samples at once. This tool is suitable for users inexperienced in RNA-seq analysis since they can leave most settings in the configuration at their default values. More experienced users will be able to change alignment parameters and so on. We welcome suggestions for additional options.

Once the configuration file is ready, the user simply types the command:

~~~
smallrnaseq -c my.conf -r
~~~

In the sections that follow, the features of the package are described all of which are accessible through the command line interface unless otherwise stated. More complex functionality is available by using the Python API and described in the online documentation.

### Counting arbitrary RNA annotations

The RNA genes in any arbitrary fasta file (referred to as ’libraries’ in the config file) may be aligned to and counts written to a csv file. Aligner indexes are first made and the names of each file are specified as a comma separated list in the order to be mapped. The normal mode of operation is to align each file only to the unmapped reads from the previous step. Alignment parameters can be set per library. This process is made clearer in Supplementary Video S1 and the documentation. This procedure is useful in determining what fraction of RNA types are present in samples. Note that miRNAs can also be counted in this manner though there is a special workflow for this as discussed in the next section. Once reads are counted the results are output to a csv file with a column for each sample. A normalized column is also included per sample. Long file names can be replaced by short labels which outputs a separate file mapping the labels to file names.

### Counting mature miRNAs

miRNA counting is simple via the command line by specifying the species in the config file. Reads are mapped to the known miRBase mature and precursor sequences directly which is a well established method used by most algorithms since it is faster then aligning to the genome. The default miRDeep2 alignment parameters for this step are ’-v 1 -a –best –strata –norc’ which we adopt as default for miRBase mapping also. Shi et al.^8^ have pointed out that miRDeep2 may tend to over count multi mappers. That is, for reads that map equally well to the positions of two or more mature miRNAs, the entire counts are added independently to the corresponding miRNA. Our default method of counting is to split the counts evenly over all mapped genes as recommended by Shi et al. The difference in results can be shown in Figure 2 which compares our counts to miRDeep2 using the default parameters for a single bovine sample file. In B with the split counts method it can be seen that certain miRNAs have higher counts for miRDeep2 (the outliers), these are genes with a number of non-unique mapping reads that are over counted e.g. bta-miR-103. This may alter results downstream for a subset of genes. The other factor that will affect quantification is the amount of padding added to mature sequences at the 5’ and 3’ ends. These can be altered in the configuration file.

**Figure 2.**
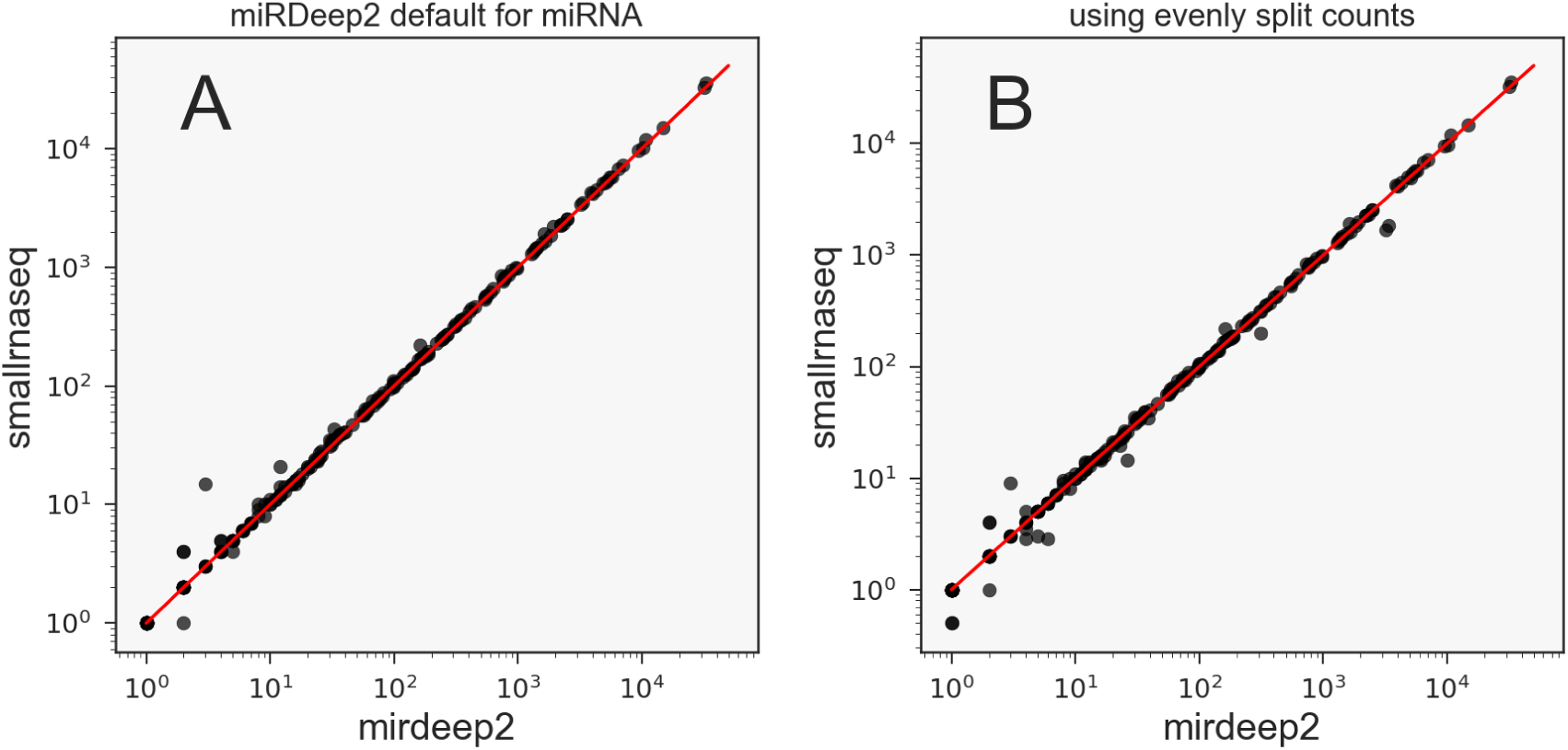
Differences in counting methods for mature miRNAs between smallrnaseq and miRDeep2 for a single sample. Several outliers below the diagonal are seen in B when we use a counting method for multi mapping reads that splits the count evenly rather than adding the totals counts equally. This indicates those genes with non unique mappings are likely to be over counted in miRDeep2. Note: outliers at bottom left of both plots are not significant.

### Counting isomiRs

The discovery of isoforms (isomiRs) that arise from the same arm as the canonical miRNA contradicts the initial view that the hairpin arm produces a single unique mature sequence. Sequence variants are common and may include 5’ and 3’ trimming and non-templated additions (enzymatically addition of a nucleotide to the 3’ end). Specific isomiRs can be substantially more abundant than the canonical form recorded in miRBase and isomiR expression appears to be tissue and population dependent^11^. miRNA expression profiling commonly uses the read counts of all isomiRs summed together. While this may be suitable in many cases it is also useful to consider the isoforms separately. There are a number of reasons isomiR counting may be a good approach:

- you only wish to count the exact canonical sequence.
- if the canonical mature sequence in miRBase is not even common in the tissue or samples you are studying and you wish to know this information.
- the dominant isoform(s) present differ from the canonical sequence such that verification with another method, e.g. a PCR assay, will fail.
- the counts of one or more isoforms can better distinguish between conditions (i.e. control and disease) than the entire sum of counts.

A single read can have more than one modification so sRNAbench^12^ provided a useful non-redundant hierarchical method of classifying isomiRs. Other naming schemes have also been used^13^. The sRNAbench scheme is used in smallrnaseq and is illustrated in Figure 3.

**Figure 3.**
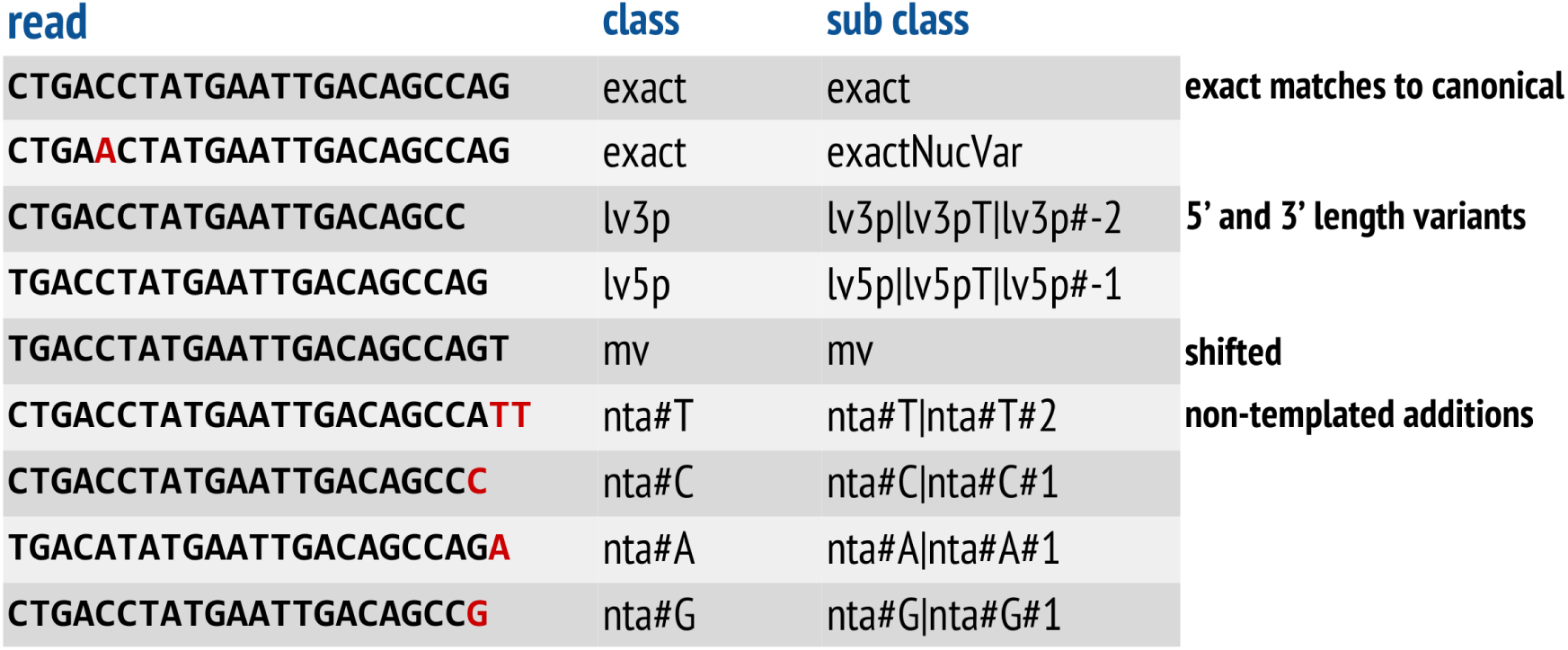
IsomiR classification scheme as used in sRNAbench.

### Novel miRNA prediction

A loose definition of a novel miRNAs is one for which the mature sequence is not present in miRBase. This may be because the species has not been well studied or insufficient evidence is available to consider a sequence a real miRNA. There are likely to be false positives in general for miRNA discovery from deep sequencing and this should be kept in mind when running an analysis of your reads. It is advised to use at least two algorithms for such an analysis. Most of the higher abundance miRNAs have been identified in the commonly studied species. Detection/prediction of new miRNAs will therefore involve looking at the low abundance (or tissue specific) forms which will need further evidence such as conservation, experimental verification and perhaps identification of function.

There are several algorithms available for predicting novel miRNAs from small RNA sequencing data^14^. miRDeep2 is most popular and contains a comprehensive algorithm for novel prediction which uses a statistical biogenisis based model to score candidates. smallrnaseq implements its own module for novel discovery, described in the methods section and online documentation. Briefly, smallrnaseq combines all samples together for analysis of read clusters. The prediction method corresponds more closely to the miRanalyzer approach^15^. Most importantly, results are output to a html file for interactive browsing. This is designed to facilitate the human annotation stage which is increasingly important for the rare or low abundance forms that remain to be discovered. This feature is demonstrated in Supplementary Video S1.

The score value is purely for the precursor classification. We recommend a default setting of 0.7 for the score cutoff to avoid removing too many true positives but this can be experimented with. This value was determined by estimating the false (FPR) and true (TPR) positive rates over a set of score cutoffs from 0.2 to 0.95. The result is shown in Figure 4. It can be seen that the estimated TPR starts to decrease between a score of 0.7 and 0.8. FPR decreases slowly though this will vary between sample types. Note that these FPR values are rough estimates and normally known RNA genes will be mapped to removed before novel prediction (see Methods). Stricter filters can also be selected to remove more candidates from the final results using additional checks.

**Figure 4.**
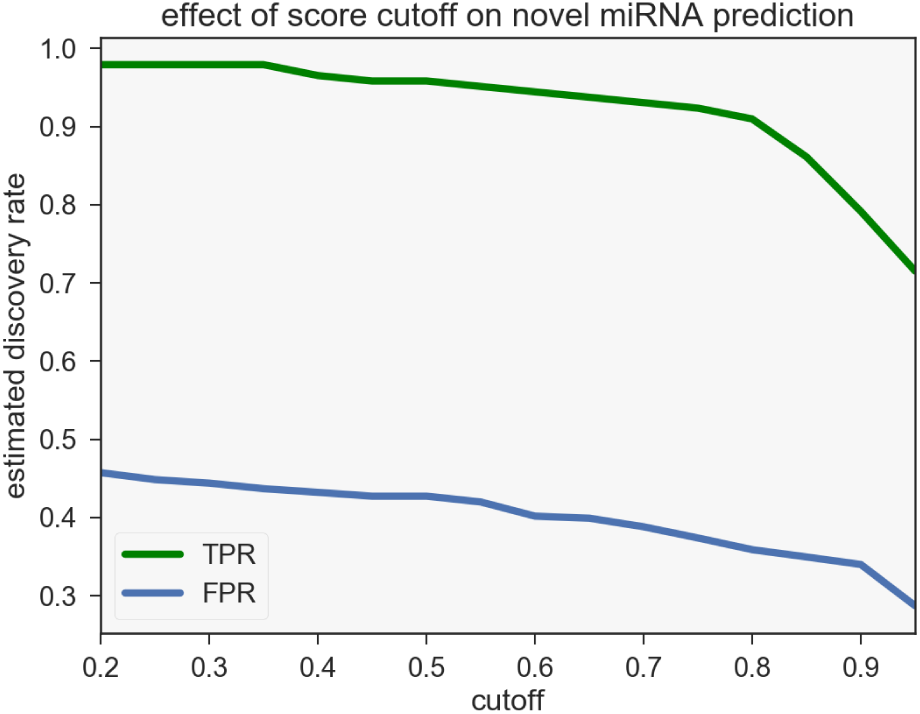
Estimates of the true positive (TPR) and false positive (FPR) novel miRNA discovery rates indicate that a score cutoff of 0.7-0.8 is suitable for general use. Details of how these values were estimated are given in Methods.

One limitation of miRDeep2 was the lack of unique identifiers for novel candidates. This leads to a many to many mapping between mature and precursor that is confusing to deal with. It also makes comparisons between independent runs difficult. Our novel discovery algorithm addresses this by providing a unique id for each mature novel candidate that always maps to the same mature sequence. Thus identical mature sequence ids will correspond between independent runs of the software. Finally our method lends itself to be scalable in that large numbers of samples can be pooled and the method run on all samples at once.

### miRDeep2 module

mirdeep2 is a popular miRNA discovery program written in perl. The command line usage is a little complicated so this module allows a configuration file to be used to provide settings. It also analyses the output results of mirdeep2 and provides summary plots.

### Counting genomic features

Counting features means counting the intersection of reads aligned to a reference genome with the locations of gene annotations (features). The features are provided typically in a gff, gtf or bed file format which are tables containing the genome coordinates and descriptor of each gene. These files are available for many species from Ensembl^16^. These will include annotations for non coding RNAs. The advantage of this approach compared to using a library is that one can map and count all features in a single step. Thus counts are normalized over all mapped genes which may be more desirable. It also allows the aligner to check for genes that map non uniquely to other locations in the genome and possibly remove them from consideration. This may be needed in the case of genes with repeat sequence or with highly conserved regions of sequence such as transfer RNAs. In the future it is intended to add a tRNA fragment counting module that requires this kind of approach^17^.

### Installation

Installation of Python packages is now relatively straightforward. Standard Python now includes the pip tool. Other dependencies are relatively minimal and can be installed easily on any linux operating system. This software is designed for use on Unix-based platforms such as linux or OS X. Windows users are recommended to install a linux operating system in a virtual machine using VirtualBox (https://www.virtualbox.org/wiki/Downloads) and run the software from there. You can access your data via shared folders. Ubuntu or Fedora based distributions of linux are recommended.

One limitation of short read aligners is that they can be memory intensive for reference genome alignment. If you are using a reference genome, which is required for novel miRNA prediction, a minimum of 8GB memory is recommended. This should especially be borne in mind if using a virtual machine where you need to allocate enough memory for the guest OS.

### Documentation

Documentation is available at https://github.com/dmnfarrell/smallrnaseq/wiki including installation instructions. Supplementary Video S1 provides an introduction and tutorial for the command line usage with two working examples. Readers are encouraged to watch this video before installing the software and using the documentation to clarify specifics.

## Discussion

Tools for sncRNA analysis from sequencing data are widely available, mostly focusing on miRNAs. As a command line tool smallrnaseq is easier to use than many others and may be attractive to users for this reason. This software is also intended to be updated regularly based on user feedback. Multiple improvements can be envisaged. Refinements to the novel miRNA discovery algorithm such as better checks for the pattern of read alignments for consistency with miRNA cleavage patterns are a priority. The process of precursor prediction will be parallelized to improve speed by using multiple threads. For this reason speed benchmarks are not detailed here. We have provided a module for assisting in differential expression analysis intended to help users do this in as few steps as possible. The method is available via the Python API and will be added to the command line interface in the next release. Also in development is a method for tRNA fragment analysis.

One of the features of scientific software in general is that they can become defunct quickly without community support^18^. We would envisage this software being potentially utilized in a number of ways that could maximize its impact. Firstly it is hoped the command line interface will find a general application. Python coders wishing to re-use the code can do so in any manner consistent with an open source license. Perhaps a more promising avenue for such tools is inclusion in community based toolkits. These provide a way to integrate disparate software into pipelines that promote best practice in analytical methods. The most relevant such project is nextgen-bcbio^19^, a framework written in Python. This provides a comprehensive set of pipelines for automated analysis of high throughput sequencing data. It is possible in the future we will integrate smallrnaseq (or some individual modules) into such a project.

## Methods

### Read counting and short read alignment

The exact method used to count reads in genes has been much discussed in the literature for RNA-seq^20^. For general users this may be confusing and a default method is selected that should give reasonable results for miRNA counting. This takes the method used in reference genome mapping of splitting non unique mapping read counts equally when adding to the mapped genes^8^. Note that the software will allow most aligners to be used by adding a method in the aligners module to wrap their mapping function. Though so far bowtie is recommended. We would be glad to integrate other aligners on request.

Read count normalization is performed by dividing counts by the total for all reads counted in the sample by default (sometimes called library normalization). There is no standard method of normalization for small RNA or miRNA that suits all cases^21^. This applies to quantitative PCR (RT-qPCR) methods as well as sequencing. Therefore other methods are available via the API when creating the count results. These will also be added to the configuration file so that the option to alter normalization method is available from the command line.

### Method for novel miRNA detection

We have implemented an approach for novel miRNA precursor prediction most closely based on the miRanalyzer (sRNAbench) method^12^, ^15^. The basic method is to find clusters of reads mapped to the genome that could be a mature sequence, extract the surrounding sequence and find suitable precursors. Structural features of each candidate precursor are then scored using a classifier. The lowest energy and highest scoring is selected.

Characteristics of the hairpin structure were encoded as features using the forgi package^22^. The features used in our algorithm are largely the same as those used in miRanalyzer and listed in Table 3 of the original paper^15^. This includes the mean free energy of the hairpin structure as calculated using RNAfold^23^. Examples of elements used as features are the length of the longest hairpin structure, number of bulges in the stem and GC-content of the loop. The miRNA hairpin structure shown in Figure 5 may help to clarify some of the terminology used. We also used the triplet-SVM features that were first proposed by Xue et al.^24^. The feature classifier was implemented with the scikit-learn library^25^ using a random forest tree predictor. Classifiers must be trained on positive and negative samples before use. A set of 2000 known human precursors were used as positives. Negatives were created by randomly extracting hairpins from known human coding sequences and filtering out obvious non-hairpins so we retain a set of ’pseudo pre-miRNAs’ as described in previous studies^24^, ^26^. The classifier can be re-trained using different data if required.

The steps for novel precursor detection are as follows:

1. reads must be mapped to a reference genome first. It is also advised to first map libraries of known RNAs to exclude them from the results first. Known miRNAs are automatically removed in this manner.
2. read counts for all samples are pooled together using each sam file and total counts for each unique read (after collapsing the original fastq files)
3. reads previously aligned to a reference genome are counted, it is also assumed the known mirbase alignments will have been removed in advance. (This is done as part of the command line workflow)
4. reads are clustered using the ClusterTrees class in bx-python
5. clusters are checked for precursors by creating multiple possible precursors in the 5’ and 3’ directions from the cluster and calculating the features of each
6. a precursor is discarded if:
  - no hairpin present
  - its read cluster overlaps with the hairpin loop
  - it has less than 19 bindings in the stem
  - it has less than 11 bindings to the region occupied by the read cluster
7. the random forest classifier/regressor is used to score the precursor features
8. the precursor with the lowest energy and highest score is used as the most likely candidate
9. the cluster region is considered the mature arm and the star sequence is estimated. If there are reads present in the star region these are added to the star count.
10. further checks on the position of the mature sequence are performed and can be used to eliminate more candidates

**Figure 5.**
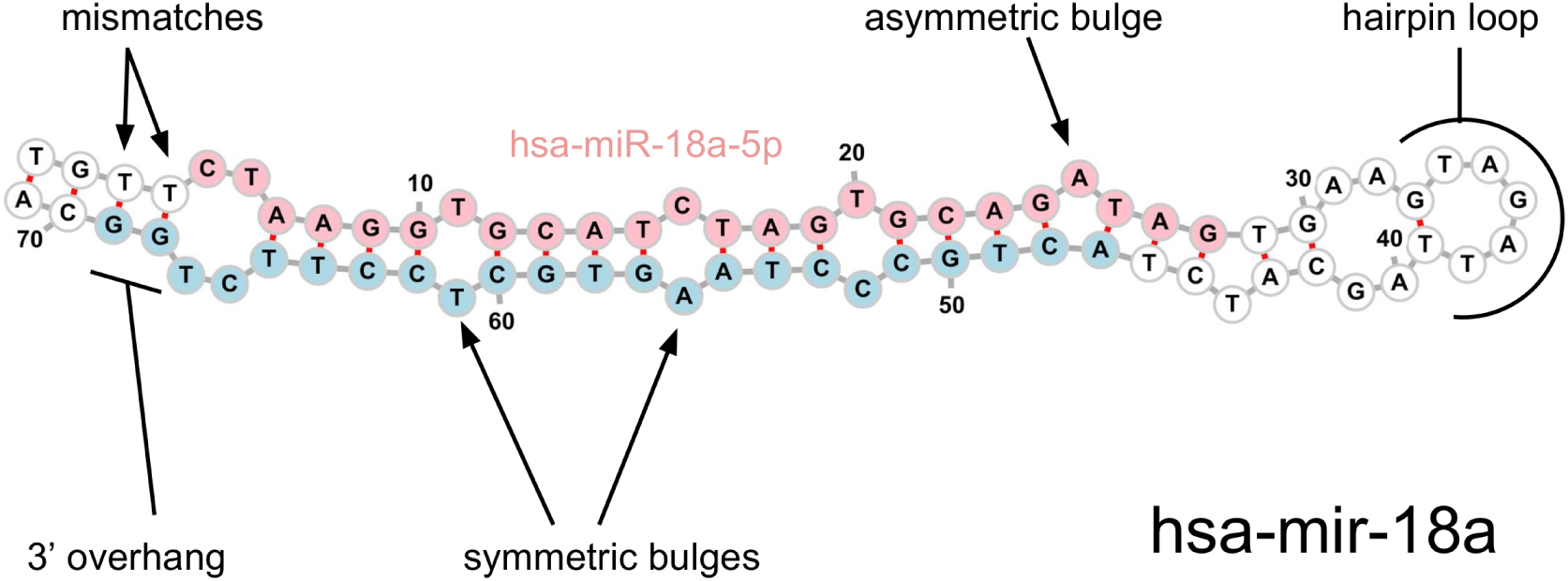
Some of the miRNA precursor features that can delineate a true miRNA. Classification algorithms can be applied to these and other features to predict likely true precursors.

The effect of score cutoffs on the novel miRNA discovery rate was examined by estimating a true positive rate (TPR) by using a single sample file. Known mature miRNAs were counted and considered to be the true positive set. Thus the TPR is the number of these miRNAs ’discovered’ in the novel prediction process as a fraction of the total positives. The process was repeated for a range of score cutoffs. A false positive rate (FPR) is somewhat harder to estimate. Since this was a bovine serum sample it is known that most false positives will be from RNA fragments in genes that might form pseudo hairpins. We therefore mapped and counted known bovine rRNA, tRNA and YRNA genes in the sample and considered these as the true negatives. The FPR is then FP/(FP+TN) at each cutoff.

### Python API and methodology

This package is entirely written in Python though uses some external software for read mapping (bowtie) and RNA folding (RNAfold). HTseq is used to assist gene counting and this relies itself on the samtools program. A flat module hierarchy is used with limited use of classes. Modules are separated broadly by function but not strictly. For example novel.py implements all the novel miRNA methods. Much of the data processing inside and between functions is handled with pandas DataFrames. This provides a very flexible table class suited to handling alignment type data without the need for a custom data structure. It is hoped that this will readily allow a Python programmer to use the core functionality without having to read a lot of documentation.

Python dependencies: Numpy, Pandas^27^, Matplotlib^28^, Seaborn^29^, HTSeq^30^, forgi^22^, bx-python^31^, pybedtools, scikit-learn^25^.

## Data Availability

This package is licensed under the Gnu Public License version v3.0. Dependencies are all freely available under various open source licenses. No proprietary software was used in this project. The github repository for this project is at https://github.com/dmnfarrell/smallrnaseq. In addition releases of the code are permanently archived on Zenodo (https://zenodo.org/record/573293). The current version is 0.2.1.

## Conclusions

The technique of next generation sequencing to quantify small non coding RNAs is a field of active research. Though there is plenty of software available they are sometimes difficult to use. Reliable standard approaches have yet to emerge in some aspects of the analysis like normalization. Software that is relatively simple but configurable is therefore appropriate. *smallrnaseq* is command line tool for processing of small RNA seq data suitable for any user. The command line interface is simple to use but extensible. Those familiar with Python may find the library useful to integrate into their workflow as a package or even standalone modules.

## Acknowledgements

D.F. is funded by an Irish Research Council Government of Ireland Postdoctoral Fellowship (GOIPD/2015/475).

## Author contributions statement

D.F. designed and wrote the software, analyzed the results and wrote the paper.

